# A comparative analysis of histone methyltransferases and demethylases in insect genome: A meta-analysis

**DOI:** 10.1101/598946

**Authors:** Parul Gulati, Surbhi Kohli, Ankita Narang, Vani Brahmachari

## Abstract

**Background:** The epigenetic regulation through post-translational modification of histones, especially methylation is well conserved, while DNA methylation is variable, being very low or absent in *Drosophila melanogaster*. Though there are several insect genomes sequenced, an analysis with a focus on their epigenetic repertoire is limited. We have compared the histone methyltransferases and the demethylases in the genome of *Drosophila melanogaster, Aedes aegypti* (Diptera), the pea aphid *Acyrthosiphon pisum*, the triatomid bug *Rhodnius prolixus* (Hemiptera), the honeybee *Apis mellifera* (Hymenoptera), the silkworm *Bombyx mori* (Lepidoptera) and the red flour beetle *Tribolium castaneum* (Coleoptera).

**Results:** We identified 38 clusters consisting of arginine, lysine methyltransferases and demethylases using OrthoFinder. To eliminate false positives, we designed a method based on identifying highly conserved domain within each class designated as the high priority domain. Out of the 9 arginine methyltransferases, *Art2, Art6* and *Art9* are identified in *D*.*melanogaster* only. We observe copy number variation between the genomes; *A*.*pisum* has nine copies of *eggless* gene (H3K9me3 methyltransferase), which can be correlated with the switch between parthenogenesis and sexual reproduction. Other than the high-priority domains, these proteins contain shared and unique domains that can mediate protein-protein interaction. Phylogenetic analysis indicates that the there is a broad conservation within the members of a class while duplication and divergence is observed in LSD1.

**Conclusion:** This meta-analysis provides a method for reliable identification of epigenetic modifiers of histones in newly sequenced insect genomes. Similar approach can be taken for other classes of genes.

## 1. Background

The post-translational modification (PTM) of histones especially at the N-terminal tail of histone is a pivotal step in epigenetic regulation during development to maintain the transcriptional status of the genes [1]. It contributes to the regulation of gene expression either by creating sites for the recruitment of specific factors or by modification of the existing sites to abolish the previous interactions [2].

The different types of modifications that exist on histones include methylation, acetylation, phosphorylation, ribosylation, succinylation, malonylation, and biotinylation [3, 4, 5, 6]. According to the “histone code hypothesis” the various post-translational modifications coexist in different combinations leading to distinct effect on gene expression [7]. The overall charge on the basic histone proteins can be changed by acetylation or phosphorylation that affects the interactions between histones and DNA. The combined effect of all the post-translational modifications brings about a change in the chromatin structure and function during development, growth, differentiation and homeostasis [8, 9]. Histone methylation is one of the well-studied PTMs, for which the essential three components, the writers, readers and the erasers have been identified [10].

Insects, the ancient group of animals which probably appeared 360 to 400 million years ago are highly diverged, occupying a prime position in the history of epigenetic phenomenon because of the diversity of polyphenisms [11]. As compared to the mammals, insects have shorter generation times, morphologically distinct development stages and high fecundity which is influenced by environmental stimulus and regulated by epigenetic mechanisms that are conserved [12]. Out of the various histone modifications known in insects, the histone methylation plays an important role; the methylated histones, as major players in the regulation of gene transcription, have been implicated in repression through heterochromatin, promoter regulation and the propagation of repressive state by DNA methylation [13].

The histone methyltransferases exhibit specificity for the histone paralogs as well as the residues they modify. The arginine methyltransferases modify histone H2A and H4 at arginine3 (R3), and H3 at positions R2, R8, R17 and R26 [14]. Lysine methyltransferases are known to modify the lysine residues at various positions in the histone H1, H2, H3 and H4 [15]. The members of protein arginine methyltransferase family catalyse the methylation of arginine in both cytoplasmic and nuclear proteins [16]. The histone H3K4, H3K36, and H3K79 methylation are known to be gene activation marks, whereas H3K9, H3K27 and H4K20 methylation gene repressive marks [17, 18]. Histone lysine methylation has important functions in biological processes like heterochromatin formation, regulation of transcription, cell cycle, genome stability and nuclear architecture [18]. It is a crucial modification that does not alter the charge of lysine residues thereby having minimal effect on the DNA-histone association. It serves as a platform for recruiting epigenetic reader-proteins which help in activating or repressing transcriptional activity [19].

Initially, methylation of histones was considered to be irreversible but in the recent years several histone demethylase families have been identified which can erase methyl marks, resulting in the reversion of the methylation effect and thus demonstrating the dynamic nature of histone methylation [20]. The histone demethylases are found in large protein complexes in association with histone deacetylases, histone methyltransferases and nuclear receptors which have an impact on the chromatin state and all chromatin-templated processes such as transcription, DNA replication, recombination and repair [21, 22]. They are involved in gene activation or repression by either actively detaching methyl group from H3K4 via the activity of its amine oxidase domain, using FAD as a cofactor [23] or demethylating H3K36 through their JmjC domain [24]. The demethylases also act on specific residues of specific histones like modifying the histone H3 at positions K4, K9, K27 and K36 [25].

In the light of the pivotal role of histone methylation, genome analysis with a focus on epigenetic tool kit is limited. We have analysed the histone methyltransferases and demethylases present in insect genomes. The representative insects considered for analysis is with the major consideration of the availability of well-annotated genome sequence and those that have been studied from the angle of their interaction with humans or as models for understanding fundamental biological process. We have considered the fruit fly *Drosophila melanogaster, Aedes aegypti* (Diptera), the pea aphid *Acyrthosiphon pisum*, the triatomid bug *Rhodnius prolixus* (Hemiptera), the honeybee *Apis mellifera* (Hymenoptera), the silkworm *Bombyx mori* (Lepidoptera) and the red flour beetle *Tribolium castaneum* (Coleoptera). *D*.*melanogaster* is a well-known model organism that has led to the discovery of several fundamental phenomena and is used to generate human disease models. *Aedes aegypti* is a vector for yellow fever, dengue, chikungunya and Zika fever [26]. *Acyrthosiphon pisum* is a sap-sucking insect and a model for the study of symbiosis, development, and host plant specialization. *Rhodnius prolixus* is the principal triatomine vector of the Chagas disease, having birds, rodents, marsupials, sloths and reptiles as host and implicated in the transmission of transposons among themselves and also to some of its vertebrate hosts, like, squirrel monkeys and opossum [27], *Apis mellifera* is the eusocial insect, has a well-structured social system and also is used in studies of pesticide toxicity, to assess non-target impact of commercial pesticides. *Bombyx mori* is an economically important insect, being a primary producer of silk. *Tribolium castaneum* is a pest particularly of food grains, and a model organism for food safety research. We have focused on histone methyltransferases and demethylases, as the number of genes assigned for this function is large among the various epigenetic players. We take into consideration that the study is dictated by the annotation of the genomes that is available in the public domain. In the present analysis we have designed an approach to identify the histone methyltransferases and demethylases in the genome with high confidence. We identified high-priority domains within each class that lead to exclusion of false positives and the identification of potential novel genes involved in epigenetic regulation. We also identified the functional domains in addition to the essential domains.

## 2. Materials and Methods

### Identification of orthogroups in the proteome

All the protein sequences from *Drosophila melanogaster, Aedes aegypti, Acyrthosiphon pisum, Rhodnius prolixus, Apis mellifera, Bombyx mori* and *Tribolium castaneum* were retrieved from UniProtKB [28]. OrthoFinder (version 2.2.6) [29] was used to identify cluster of orthologous genes. To generate a non-redundant list of Uniprot IDs for histone methyltransferases (arginine and lysine) and demethylases from Drosophila the following sources were used: keyword based literature search, the Flybase and Uniprot. This list was used to identify the relevant orthoclusters for the three classes of proteins of the selected insect genomes. We also mapped duplications in selected orthogroups or orthoclusters. We devised an additional approach to mine novel genes in these classes that are designated either as hypothetical or uncharacterised proteins. Interproscan [30] was used to identify the conserved domains for each gene.

### Phylogenetic analysis of curated orthogroups

The inferred amino acid sequences of each of the arginine, lysine methyltransferases and the demethylases were aligned separately with MAFFT version 7 L-INS-i [31]. The data set comprised of 49 arginine, 115 lysine methyltransferases and 100 demethylases. Alignment was obtained in the CLUSTAL format. Phylogenetic trees were drawn with these alignments with default parameters. Tree file without terminal node number in the Newick format was used for drawing tree and colour coded using the software Rainbow Tree [32, 33]. The whole proteome based species tree generated by OrthoFinder was used for comparison. The complete workflow for the analysis is given in Figure 1.

**Figure 1.**
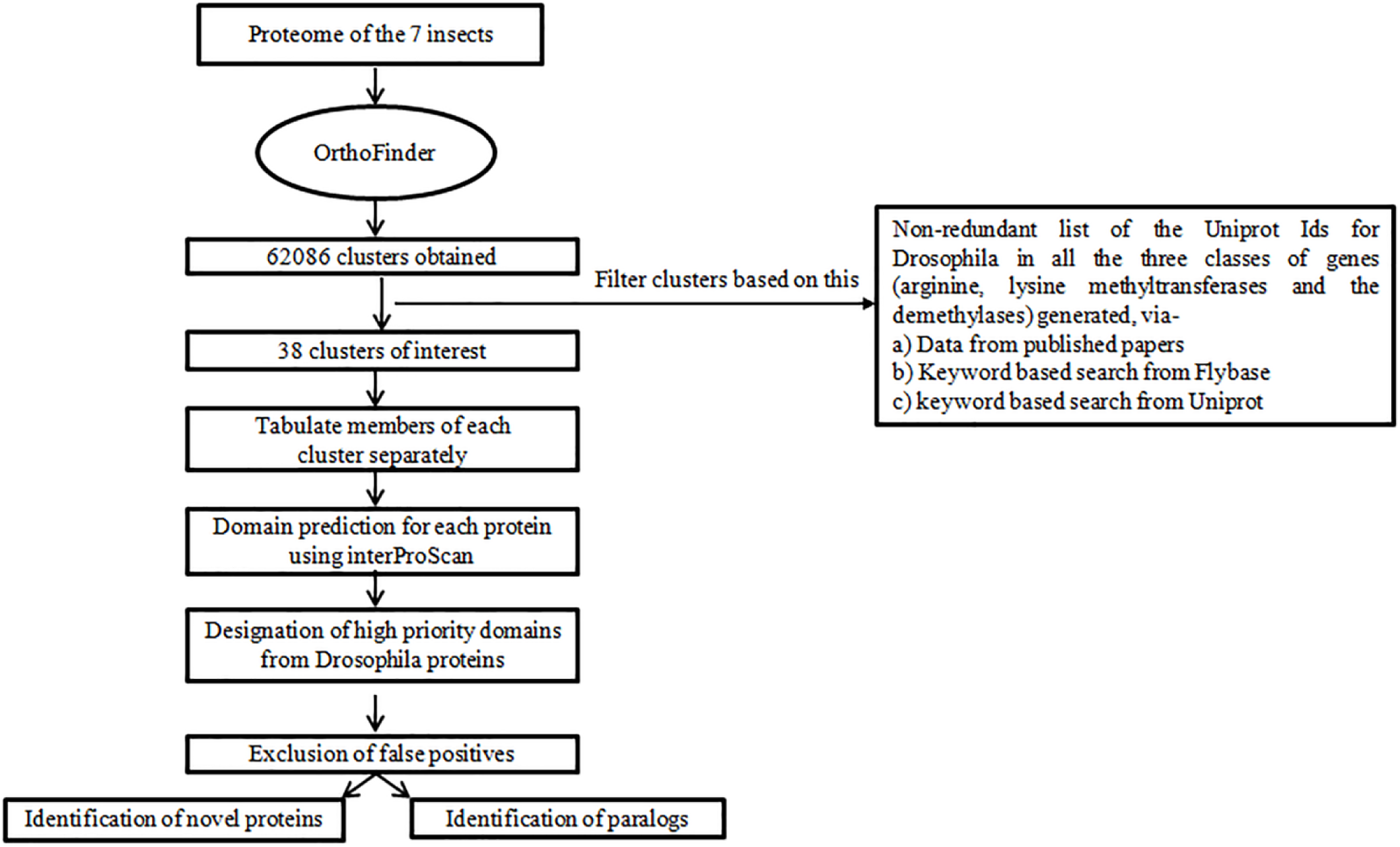
Workflow followed for the *in silico* analysis of histone methyltransferases and demethylases

## 3. Results and Discussion

### Retrieval of curated methylases and demethylases orthoclusters

From OrthoFinder we obtained 62086 Orthoclusters. Using Uniprot IDs and keyword based searches 38 clusters for histone arginine, lysine methyltransferases and the demethylases were identified by manual curation (Table 1). The filtered data comprised 16 clusters of lysine methyltransferases and 10 clusters of the nine arginine methyltransferases in Drosophila.

**Table 1:**
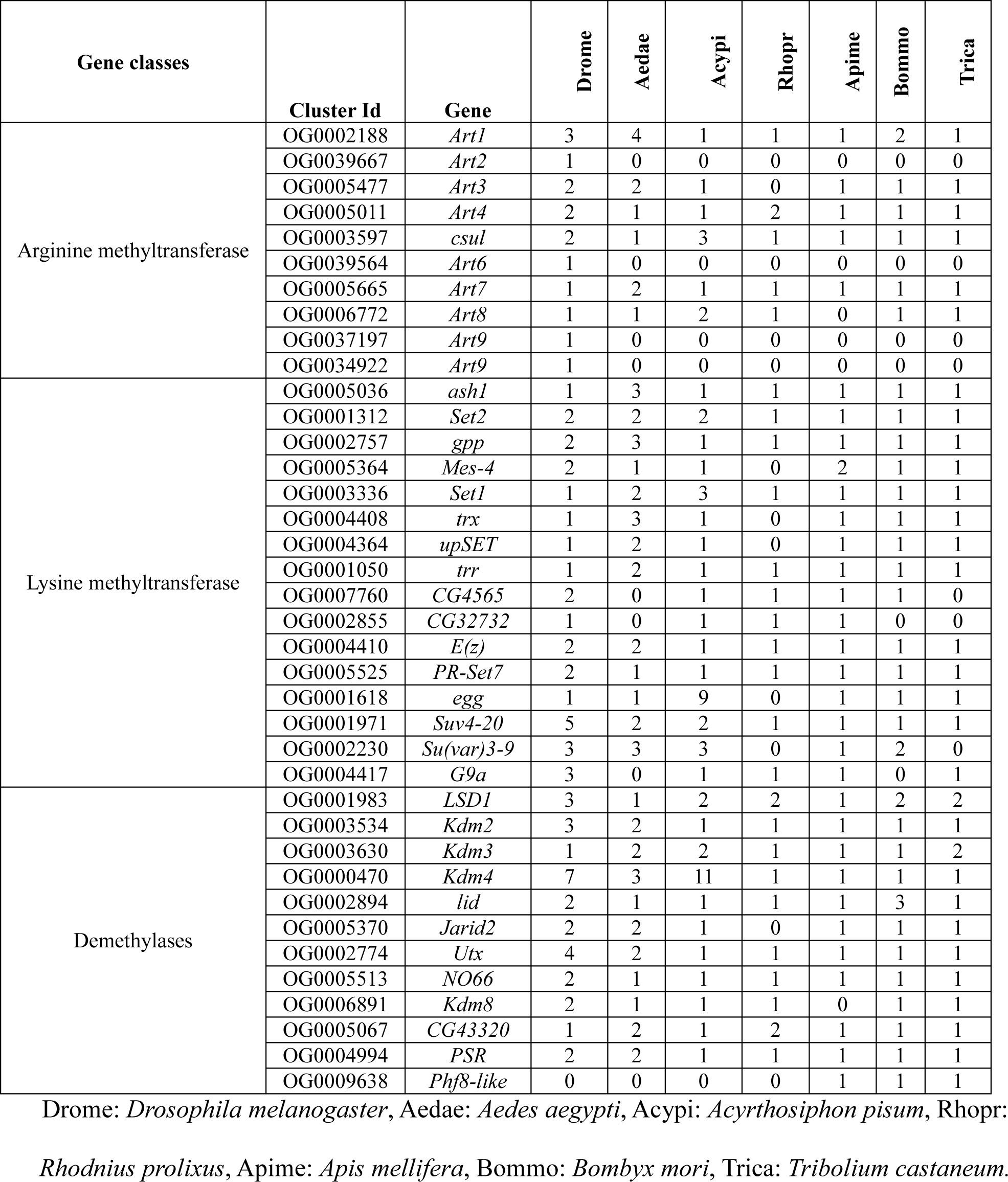
The clusters for histone methyltransferases and the demethylases in the various representative species after curation along with the number of homologs for each insect in the cluster

| Gene classes | Cluster Id | Gene | Drome | Aedae | Acypi | Rhopr | Apime | Bommo | Trica |
| --- | --- | --- | --- | --- | --- | --- | --- | --- | --- |
| Arginine methyltransferase | OG0002188 | *Art1* | 3 | 4 | 1 | 1 | 1 | 2 | 1 |
|  | OG0039667 | *Art2* | 1 | 0 | 0 | 0 | 0 | 0 | 0 |
|  | OG0005477 | *Art3* | 2 | 2 | 1 | 0 | 1 | 1 | 1 |
|  | OG0005011 | *Art4* | 2 | 1 | 1 | 2 | 1 | 1 | 1 |
|  | OG0003597 | *csul* | 2 | 1 | 3 | 1 | 1 | 1 | 1 |
|  | OG0039564 | *Art6* | 1 | 0 | 0 | 0 | 0 | 0 | 0 |
|  | OG0005665 | *Art7* | 1 | 2 | 1 | 1 | 1 | 1 | 1 |
|  | OG0006772 | *Art8* | 1 | 1 | 2 | 1 | 0 | 1 | 1 |
|  | OG0037197 | *Art9* | 1 | 0 | 0 | 0 | 0 | 0 | 0 |
|  | OG0034922 | *Art9* | 1 | 0 | 0 | 0 | 0 | 0 | 0 |
| Lysine methyltransferase | OG0005036 | *ash1* | 1 | 3 | 1 | 1 | 1 | 1 | 1 |
|  | OG0001312 | *Set2* | 2 | 2 | 2 | 1 | 1 | 1 | 1 |
|  | OG0002757 | *gpp* | 2 | 3 | 1 | 1 | 1 | 1 | 1 |
|  | OG0005364 | *Mes-4* | 2 | 1 | 1 | 0 | 2 | 1 | 1 |
|  | OG0003336 | *Set1* | 1 | 2 | 3 | 1 | 1 | 1 | 1 |
|  | OG0004408 | *trx* | 1 | 3 | 1 | 0 | 1 | 1 | 1 |
|  | OG0004364 | *upSET* | 1 | 2 | 1 | 0 | 1 | 1 | 1 |
|  | OG0001050 | *trr* | 1 | 2 | 1 | 1 | 1 | 1 | 1 |
|  | OG0007760 | *CG4565* | 2 | 0 | 1 | 1 | 1 | 1 | 0 |
|  | OG0002855 | *CG32732* | 1 | 0 | 1 | 1 | 1 | 0 | 0 |
|  | OG0004410 | *E(z)* | 2 | 2 | 1 | 1 | 1 | 1 | 1 |
|  | OG0005525 | *PR-Set7* | 2 | 1 | 1 | 1 | 1 | 1 | 1 |
|  | OG0001618 | *egg* | 1 | 1 | 9 | 0 | 1 | 1 | 1 |
|  | OG0001971 | *Suv4-20* | 5 | 2 | 2 | 1 | 1 | 1 | 1 |
|  | OG0002230 | *Su(var)3-9* | 3 | 3 | 3 | 0 | 1 | 2 | 0 |
|  | OG0004417 | *G9a* | 3 | 0 | 1 | 1 | 1 | 0 | 1 |
| Demethylases | OG0001983 | *LSD1* | 3 | 1 | 2 | 2 | 1 | 2 | 2 |
|  | OG0003534 | *Kdm2* | 3 | 2 | 1 | 1 | 1 | 1 | 1 |
|  | OG0003630 | *Kdm3* | 1 | 2 | 2 | 1 | 1 | 1 | 2 |
|  | OG0000470 | *Kdm4* | 7 | 3 | 11 | 1 | 1 | 1 | 1 |
|  | OG0002894 | *lid* | 2 | 1 | 1 | 1 | 1 | 3 | 1 |
|  | OG0005370 | *Jarid2* | 2 | 2 | 1 | 0 | 1 | 1 | 1 |
|  | OG0002774 | *Utx* | 4 | 2 | 1 | 1 | 1 | 1 | 1 |
|  | OG0005513 | *NO66* | 2 | 1 | 1 | 1 | 1 | 1 | 1 |
|  | OG0006891 | *Kdm8* | 2 | 1 | 1 | 1 | 0 | 1 | 1 |
|  | OG0005067 | *CG43320* | 1 | 2 | 1 | 2 | 1 | 1 | 1 |
|  | OG0004994 | *PSR* | 2 | 2 | 1 | 1 | 1 | 1 | 1 |
|  | OG0009638 | *Phf8-like* | 0 | 0 | 0 | 0 | 1 | 1 | 1 |

Drome: *Drosophila melanogaster*, Aedae: *Aedes aegypti*, Acypi: *Acyrtosiphon pisum*, Rhopr:
*Rhodnius prolixus*, Apime: *Apis mellifera*, Bommo: *Bombyx mori*, Trica: *Tribolium castaneum*.

*Art9* is identified only in *D*. *melanogaster* and the two isoforms (99.2% identity), were grouped into two singleton clusters as additional 53 amino acids are present at the N-terminal in one of them. As against the nine arginine methyltransferases in Drosophila, only six were identified in Aedes, Acyrthosiphon, Bombyx and Tribolium while only five arginine methyltransferases were identified in Rhodnius and Apis genome. *Art2, Art6* and *Art9* methyltransferases are present in *D*.*melanogaster* but were absent in all other insects. No redundancy in activity has been reported for the methyltransferases with respect to the position of methylation at various residues and human arginine methyltransferases (PRMT) have non-overlapping properties and are thought to be involved in different cellular processes [34].

For lysine methyltransferases (KMT) Drosophila, Aedes and Apis contained all 16 KMTs while two or more clusters were absent in all other insects. Rhodnius genes were represented in only eleven clusters. The lysine methyltransferases do show redundancy as clusters comprising the *Set2, CG4565* and *CG32732* enzymes are responsible for methylating H3 at K36 with *CG32732* as the only protein which has both H3K4 and H3K36 methylation activity [35]. RNA interference-mediated (RNAi) suppression of Drosophila *Set2* leads to the lack H3K36 methylation, suggesting its crucial role in depositing this activating modification [36].

The histone demethylases were represented by 12 clusters. The *Kdm4* (lysine demethylase) clusters show co-existence of the closely related members-*Kdm4a* and *Kdm4b* genes which are essential for mediating the ecdysteroid hormone signalling and are biologically redundant [37]. The trend shown by the cluster of methyltransferases is also followed in this class with Rhodnius proteins represented in least number of clusters.

The clustering in OrthoFinder is based on the identity in sequence of amino acids. However this may not represent the presence of the relevant functional domain in the protein. Therefore the data was curated to exclude the false positive members from the clusters based on the presence of certain domains. The reliable signature for a given functional class was identified in Drosophila proteins as High-priority domains. The domain(s) that was present at high frequency in each class of protein was identified (Figure 2). For example, S-adenosyl-L-methionine-dependent methyltransferase domain is present in 9/9 genes in Drosophila, while PH domain like is present in 2/9 cases. Thus we selected the SAM-dependent MTase domain as the high priority domain for Arginine methyltransferase while the SET domain with or without the Pre-SET or Post-SET domain is the high priority domain for the lysine methyltransferases. Demethylases have the JmjC and/or the JmjN as the high priority domain. Based on this criterion, the output of OrthoFinder was filtered and the overall false positive rate was observed to be 8.2%. On the same criteria 37% of the members in the clusters qualify as putative methyltransferases or demethylases. This is a stringent criterion and there is a chance that a putative methyltransferase gene may get excluded, as 8 false positives were identified in Drosophila. While this can be due to partial sequence being present in the database, the genes identified based on this criterion could be a confirmed methyltransferase/demethylase. False positive cases for different insect proteomes are provided in Additional File 1 A. While analysing the proteins further, we observed that only 54.51 % of the proteins have well defined functions (known or already annotated proteins) while 37.29 % are novel which are annotated either as uncharacterized or hypothetical proteins (Additional File 1 B). Thus by this exercise we have been able to functionally classify a considerable number of proteins previously termed as uncharacterized.

**Figure 2.**
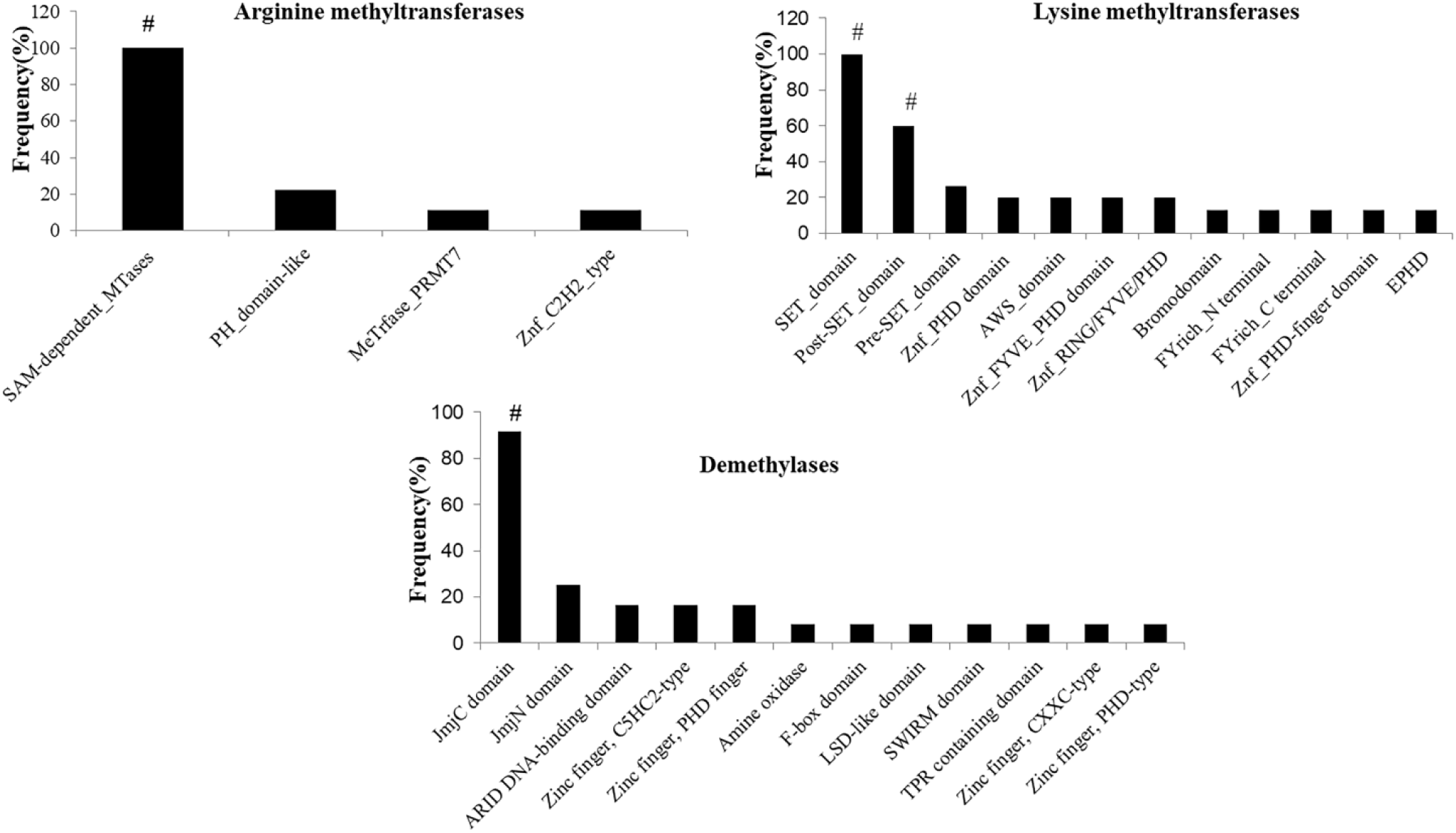
The frequency of occurrence of the domains in Drosophila. The three classes of enzymes used for domain identification are shown in the figure. The percentage of occurrence of each domain is plotted on the Y-axis, # - high priority domains.

We also observed paralogs, described as the same genes/proteins within the genome of a species, derived from a common ancestor gene through duplication events. *A*.*pisum* and *A*.*aegypti* showed maximum number of paralogs while rest of the insects show a little or no duplication in the methyltransferases and the demethylases (Table 2). *A*.*pisum* genome is known to be duplicated for the chromatin modifiers [38]. The highest number of paralogs appearing in *A*.*pisum* corresponds to the Kdm4a/b and the *eggless*/*SETDB1*gene*s* having 11 and 9 copies respectively.

**Table 2:** The number of paralogs for the methyltransferase and demethylase genes in the various representative species.

| Orthocluster | Gene | Drome | Aedae | Acypi | Rhopr | Apime | Bommo | Trica |
| --- | --- | --- | --- | --- | --- | --- | --- | --- |
| OG0002188 | *Art1* | 0 | 2 | 0 | 0 | 0 | 0 | 0 |
| OG0005011 | *Art4* | 0 | 0 | 0 | 2 | 0 | 0 | 0 |
| OG0003597 | *csul* | 0 | 0 | 3 | 0 | 0 | 0 | 0 |
| OG0005665 | *Art7* | 0 | 2 | 0 | 0 | 0 | 0 | 0 |
| OG0006772 | *Art8* | 0 | 0 | 2 | 0 | - | 0 | 0 |
| OG0001312 | *Set2* | 0 | 0 | 2 | 0 | 0 | 0 | 0 |
| OG0003336 | *Set1* | 0 | 0 | 3 | 0 | 0 | 0 | 0 |
| OG0004408 | *trx* | 0 | 2 | 0 | - | 0 | 0 | 0 |
| OG0004410 | *E(z)* | 0 | 2 | 0 | 0 | 0 | 0 | 0 |
| OG0001618 | *egg* | 0 | 0 | 9 | - | 0 | 0 | 0 |
| OG0001971 | *Suv4-20* | 0 | 0 | 2 | 0 | 0 | 0 | 0 |
| OG0002230 | *Su(var)3-9* | 0 | 0 | 3 | - | 0 | 0 | - |
| OG0001983 | *LSD1* | 0 | 0 | 2 | 2 | 0 | 2 | 2 |
| OG0003630 | *Kdm3* | 0 | 2 | 2 | 0 | 0 | 0 | 0 |
| OG0000470 | *Kdm4* | 2 | 2 | 11 | 0 | 0 | 0 | 0 |
| OG0002894 | *lid* | 0 | 0 | 0 | 0 | 0 | 2 | 0 |
| OG0002774 | *Utx* | 0 | 2 | 0 | 0 | 0 | 0 | 0 |
| OG0005067 | *CG43320* | 0 | 2 | 0 | 0 | 0 | 0 | 0 |
| OG0004994 | *PSR* | 0 | 2 | 0 | 0 | 0 | 0 | 0 |

The numerals indicate the number, 0 represents absence of paralogs (single copy of the gene present) and – the absence of the homolog. Drome: *Drosophila melanogaster*, Aedae: *Aedes aegypti*, Acypi: *Acyrtosiphon pisum*, Rhopr: *Rhodnius prolixus*, Apime: *Apis mellifera*, Bommo: *Bombyx mori*, Trica: *Tribolium castaneum*.

The Drosophila eggless/dSetdb1 protein is responsible for trimethylation of H3K9 and is essential for viability and fertility [39, 40]. It is required at multiple stages of oogenesis, maintenance and differentiation of Germline and Follicle Stem Cells. It is involved in piRNA cluster transcription and Dpp (Decapentaplegic) signalling during oogenesis, expression of specific long non-coding RNAs, apoptosis-related gene regulation, and silencing of key spermatogenesis gene *Phf7* [41, 42, 43]. The gene *eggless* along with the *sxl* (sex determining) gene preserves the female fate by conferring the repressive mark H3K9me3 on *Phf7* gene, a histone reader that associates with H3K4me2 and pilots the male sexual program in the germ line [44]. *A*.*pisum* undergoes cyclic parthenogenesis (10-30 generations) followed by a single sexual cycle. The switch between these two modes of reproduction is unclear but considering the role of Setdb1 in maintaining female identity its involvement in parthenogenesis cannot be ruled out. Therefore the presence of high number of *eggless* in *A*.*pisum* could be associated with the switching of the mode of reproduction. The switch is also known to be sensitive to environmental signals like photoperiod and temperature [45]. It is interesting to speculate that epigenetic alteration leading to altered gene expression forms an interphase between environment and the genotype.

The Kdm4 family of demethylases is highly conserved. It removes di- and tri-methyl groups from H3K9 and H3K36 in Drosophila, *C*.*elegans*, and mice. Drosophila KDM4 is required for maintaining the normal structure and function of heterochromatin, essential for spatial arrangement of repetitive elements, and is involved in Position effect variegation (PEV). It is important for double strand break (DSB) movement in heterochromatin and its loss leads to a delay in DSB repair and an increase in homologous recombination (HR) repair at heterochromatic DSBs. KDM4 specifically promotes demethylation of H3K9me3 and H3K56me3 at heterochromatic DSBs [46, 47, 48]. The biologically redundant members KDM4a and KDM4b are essential for ecdysteroid hormone signalling in Drosophila [37]. The loss of KDM4 leads to development arrest due to increased silencing marks at H3K9me2, 3 and the transcriptional activation of ecdysone response genes [37]. Increased titres of ecdysone in *A*.*pisum* leads to activation of Br-C complex and ultimately contributes to formation of wingless offspring. As wingless is more favoured phenotype under normal conditions as opposed to stressed conditions (eg aphids high density) where asexual adult females form winged morphs instead of wingless by altering the developmental fate of the embryos [49], high copy number of *Kdm4* could be correlated to wing polyphenism. The above two examples also show that epigenetics is present at the interface of environment and genotype interaction.

### Domain architecture

We analysed the occurrence and sharing of domains other than the high-priority domains in a given methyltransferase/ demethylase in the different genomes (Table 3). The domains involved in protein-protein interaction like the bromodomain, Zinc finger domain, SWIRM domain are present in both lysine methyltransferases and demethylases, though not all the domains are shared by the genes in all the genomes we have analysed. The PRMT5 oligomerization domain is present in PRMT5 of the genomes analysed. Art8 of Rhodnius has methyltransferase Fkbm domain. The function of this domain is unknown in Rhodnius, but the known function of such a domain in Streptomyces is in a specific methylation step in the biosynthesis of the immunosuppressant. One can speculate similar function of reduction of inflammation in the host during its blood meal, as known for the bioactive molecules in the saliva of Rhodnius [50]. Another domain we find only in Ash1 of Rhodnius is the Ubiquitin system component Cue domain which is involved in the binding of ubiquitin conjugating enzymes to epigenetic complexes. The Set2Rpb1 interacting domain is found in Set2 of several genomes and is implicated in coupling histone H3 K36 methylation with transcription elongation [51]. The presence of RNA recognition domain in *Set1/SetD1* gene in most of the genomes analysed is relevant as it marks transcription start site [52].

**Table 3:**
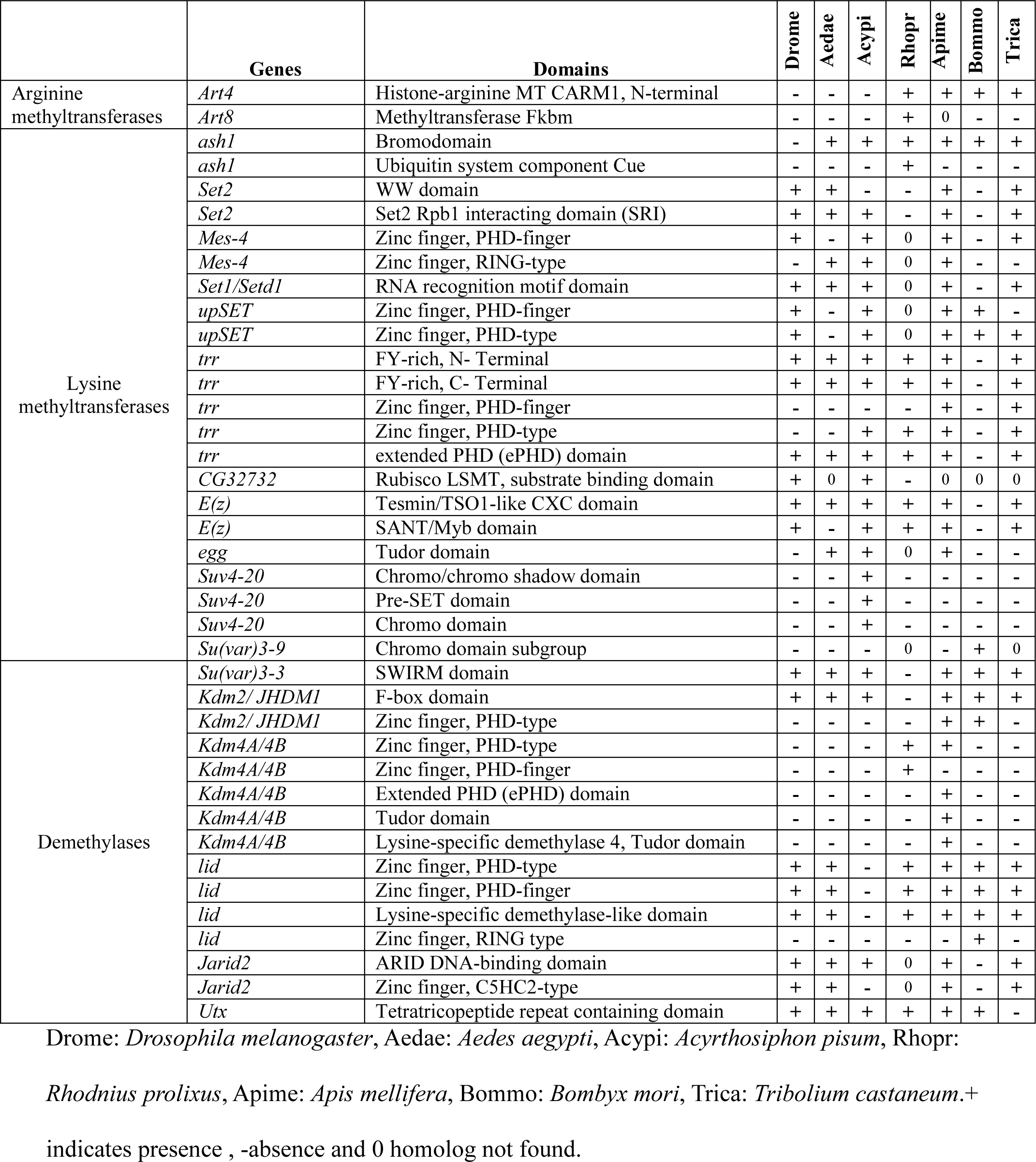
The conservation of various functional domains in the methyltransferases and the demethylases. The domain names are taken from Interproscan.

The methyltransferases and demethylases have acquired domains (in addition to the catalytic domains), that are important for interactions either with other proteins or with RNA/DNA. This will facilitate the formation of different complexes in various combinations to generate complexes of unique composition that can be targeted to specific sites to bring about the repression or activation through epigenetic modifications (Maini et al. under review). In addition, they may interact with proteins relevant for their interaction with environmental signals depending upon their habitat.

### Evolutionary relationships

On comparison with total proteome phylogram (Additional File 2), arginine methyltransferases, lysine methylatransferases and demethylases showed variation in terms of evolutionary relationships among insect species of different orders. The species belonging to the same order showed high conservation among themselves than those belonging to different orders with some exceptions.

Although for majority of arginine methyltransferases of the dipterans, Drosophila and Aedes showed more conservation among themselves than that of other insects, Art3 and Art1 were exceptions, suggesting their variability among the arginine methyltransferases (Figure 3). On the other hand Acyrthosiphon and Rhodnius belonging to same order show less conservation suggesting the epigenetic tool kit though conserved can still accumulate species specific variation. Lysine methyltransferases showed considerable deviation from the phylogenetic tree derived from the complete proteome (Figure 4). On analysing each protein it was interesting to note that even species of the same order like Diptera (Drosophila and Aedes) and Hemiptera (Acyrthosiphon and Rhodnius) branched together only 25% and 12.5% times respectively. These finding suggests that lysine methyltransferases are more species specific.

**Figure 3.**
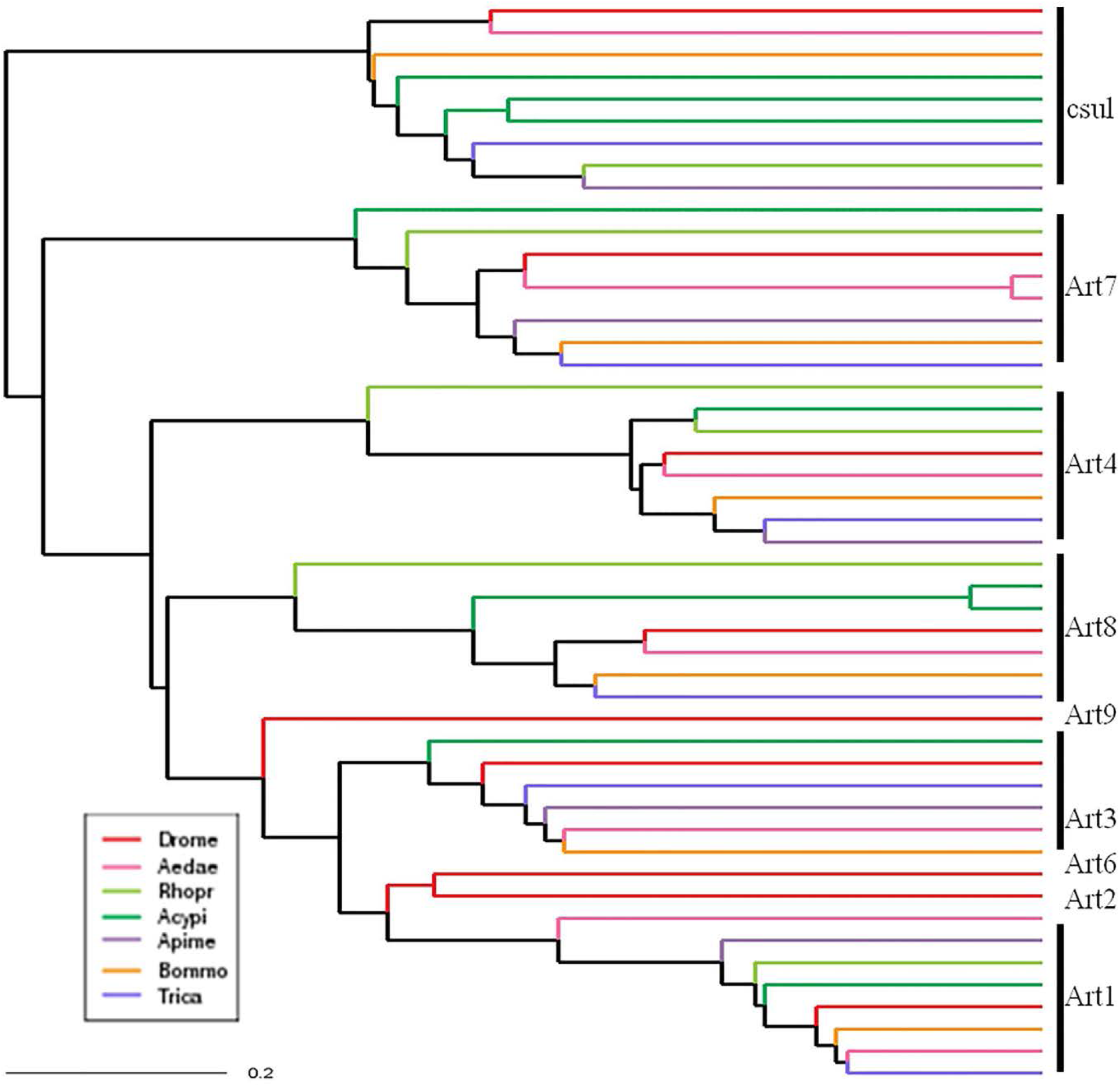
Phylogenetic tree for the arginine methyltransferases. The coloured lines indicate proteins from different insects as specified in the inset. Drome: *Drosophila melanogaster*, Aedae: *Aedes aegypti*, Acypi: *Acyrthosiphon pisum*, Rhopr: *Rhodnius prolixus*, Apime: *Apis mellifera*, Bommo: *Bombyx mori*, Trica: *Tribolium castaneum*.

**Figure 4.**
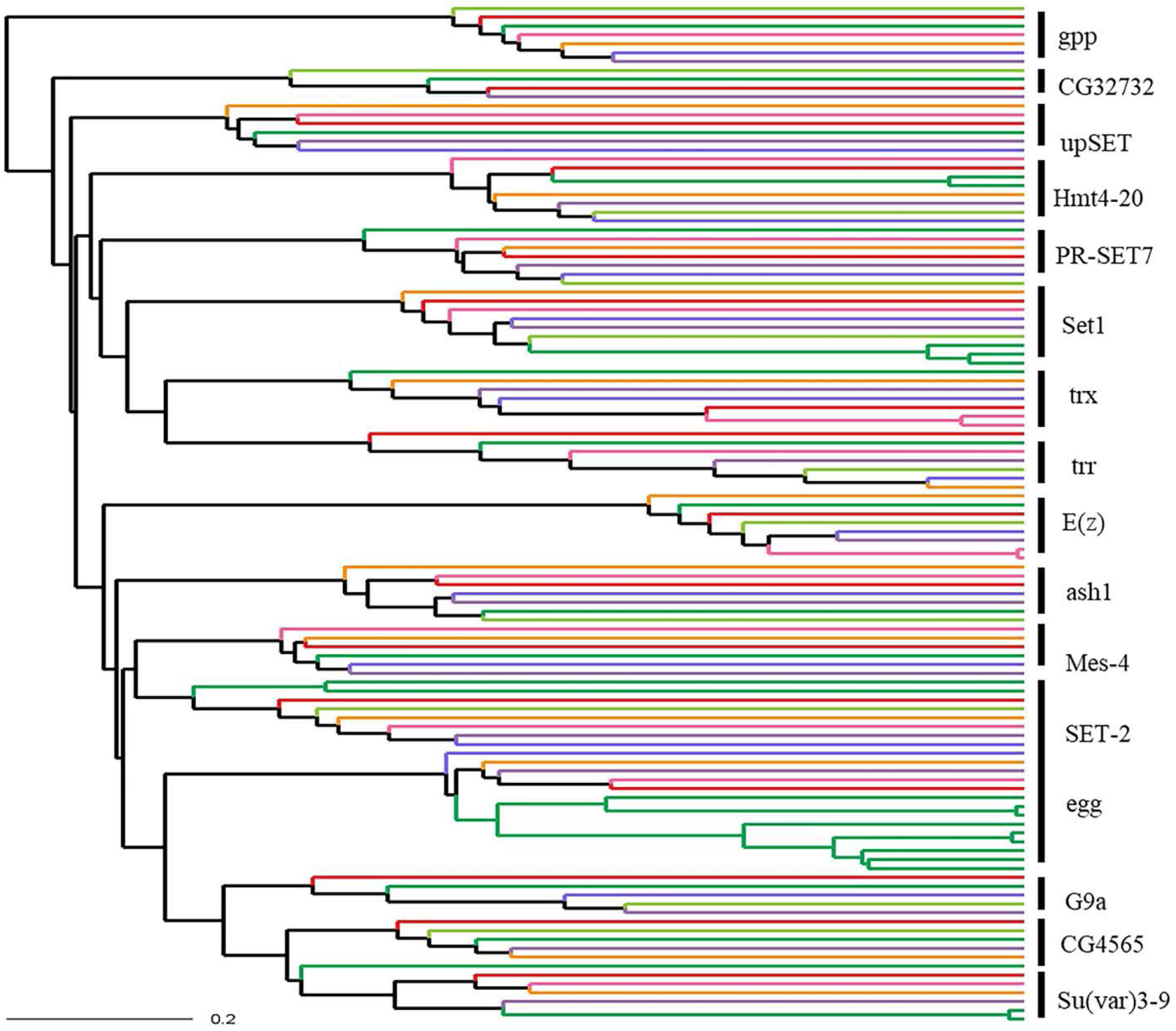
Phylogenetic tree for the lysine methyltransferases. The coloured lines indicate proteins from different insects as specified in the inset. Drome: *Drosophila melanogaster*, Aedae: *Aedes aegypti*, Acypi: *Acyrthosiphon pisum*, Rhopr: *Rhodnius prolixus*, Apime: *Apis mellifera*, Bommo: *Bombyx mori*, Trica: *Tribolium castaneum*

Similarly histone demethylases were highly variable than complete proteome phylogenetic tree with some proteins showing more conservation in terms of insect orders than others. One interesting example among demethylases is LSD1, the only amine oxidase demethylase in insects which had multiple homologs present in a given organism which show divergence, thus suggesting a duplication event followed by divergence (Figure 5). In Acyrthosiphon, Rhodnius, Bombyx and Tribolium except Dipterans (Drosophila, Aedes) and Hymenopterans (Apis) two copies of this protein diverge into two branches. This could either represent a loss of protein in these orders or duplication of the same in other insects. Duplication and divergence of this protein could reflect their tissue specific or developmental stage specific expression or divergence leading to non-histone substrate targeting or the variation in the interaction with environmental signals.

**Figure 5.**
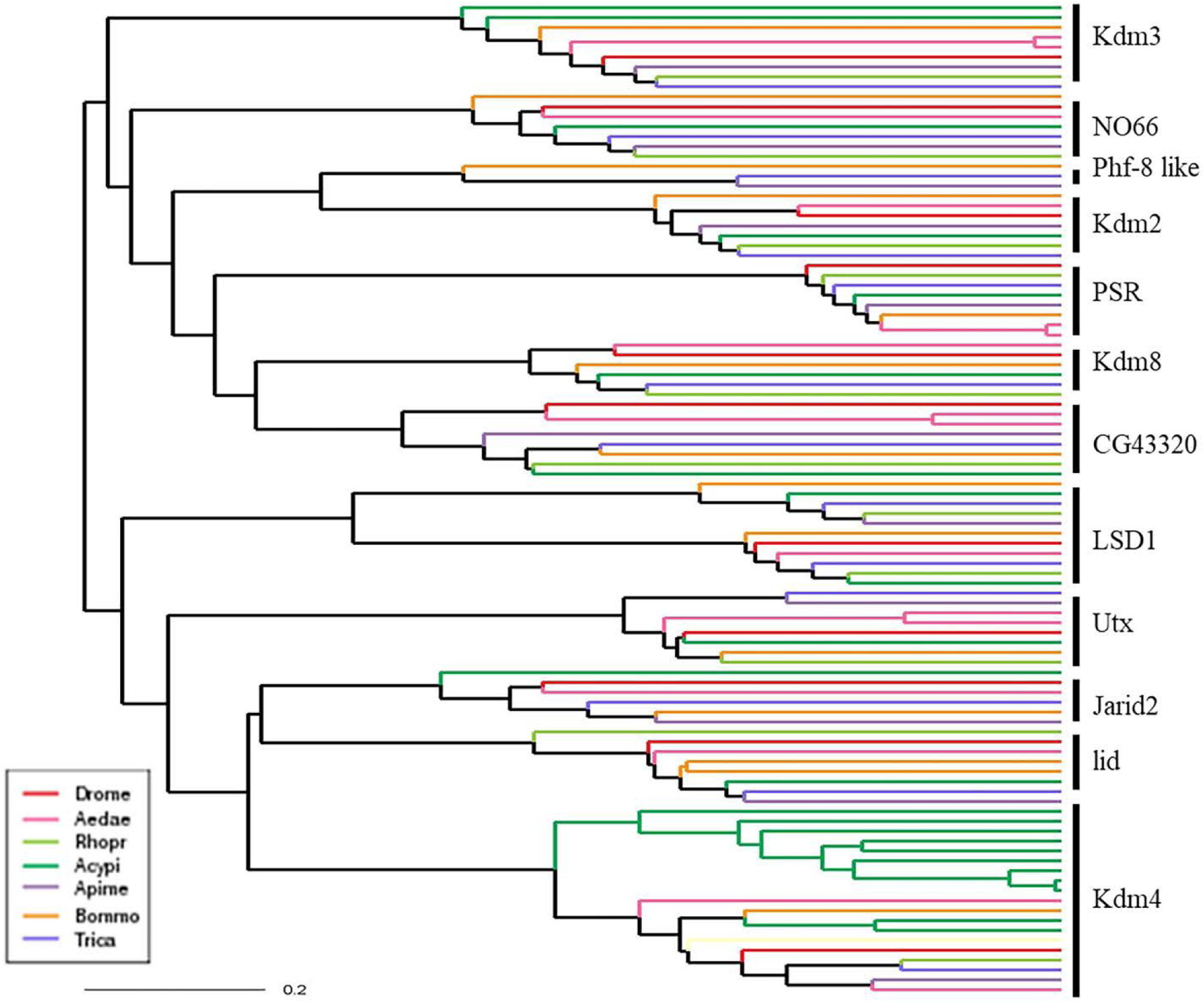
Phylogenetic tree for demethylases. The coloured lines indicate proteins from different insects as specified in the inset. Drome: *Drosophila melanogaster*, Aedae: *Aedes aegypti*, Acypi: *Acyrthosiphon pisum*, Rhopr: Rhodnius *prolixus*, Apime: *Apis mellifera*, Bommo: *Bombyx mori*, Trica: *Tribolium castaneum*.

## 4. Summary

The protein coding capacity of the insect genomes appears to be inadequate to achieve the complexity of these systems. One of the ways in which this is compensated can be attributed to globally acting epigenetic modifiers. This machinery can utilize limited repertoire of protein coding genes in different combinations and various contexts, in addition to the regulation by non-coding RNA, to achieve unique outcomes. The epigenetic regulation is recognized as an important player in maintaining this diversity of functions. Here, we analysed the genomes of the insects of order Diptera, Hemiptera, Hymenoptera, Lepidoptera and Coleoptera for the epigenetic modifiers. Histone modifications play crucial role in maintaining epigenetic status of an organism; the most common role played by the methyltransferases. The proteins having different domains can be implied in functions other than histone methylation, reflecting the additional functions that they may carry out as perhaps moonlighting activity.

The diversity in the domain architecture therefore could be a reflection of gain/loss of additional functions, also contribute to their participation in different functional complexes. Thus epigenetic modifiers are important contributors to the economy of the genomes with reference to their coding capacity. The current analysis will serve as a resource for mining epigenetic modifiers from whole genome data.

## Declarations

### Ethics approval and consent to participate

Not Applicable

## Consent for publication

Not Applicable

## Availability of data and material

All data generated or analysed during this study are included in this published article [and Additional files].

## Authors’ contributions

PG and SK-carried out the work, analysed the results and writing the manuscript, AN-Cluster generation using OrthoFinder, VB-conceptualization of the work, analysis of the results and writing the manuscript.

## Abbreviations

KMT: lysine methyltransferases
PRMT: Protein arginine methyltransferases
PTM: post translational modification
DSB: double strand break

## Acknowledgements

The authors gratefully acknowledge the funding for the work through research grant to VB from the Council for Scientific and Industrial Research (CSIR), (EpiHeD: BSC0118/2012-17) and SERB No.60 (0102)/12/EMR-II), Government of India. We acknowledge DBT Bioinformatics facility at ACBR and the UGC SAP-II support. PG acknowledges CSIR, SK and AN acknowledge University Grants Commission, Government of India for support through Senior Research Fellowship and D.S. Kothari Post-doctoral fellowship

## Additional files

Gulati et al Additional File 1.docx **Additional File 1:** Different Insect proteomes showing A. False positive rates B: Novel proteins

Gulati et al Additional File 2.tiff **Additional File 2.** Phylogenetic tree obtained from OrthoFinder for the whole proteome of the seven species used in the study. Drome: *Drosophila melanogaster*, Aedae: *Aedes aegypti*, Acypi: *Acyrthosiphon pisum*, Rhopr: *Rhodnius prolixus*, Apime: *Apis mellifera*, Bommo: *Bombyx mori*, Trica: *Tribolium castaneum*

## References

1. Imai K and Ochiai K. Role of histone modification on transcriptional regulation and HIV-1 gene expression: possible mechanisms of periodontal diseases in AIDS progression. Journal of Oral Science 2011; 53, 1–13.

2. Khare S P, Habib F, Sharma R, Gadewal N, Gupta S and Galande S. Histome - a relational knowledgebase of human histone proteins and histone modifying enzymes Nucleic acid Research 2012; 40, D337–D342.

3. Rothbart S B, Strahl B D. Interpreting the language of histone and DNA modifications. Biochim. Biophys. Acta, Gene Regul. Mech. 2014; 1839, 627.

4. Wozniak G. G.; Strahl B. D. Hitting the ’mark’: interpreting lysine methylation in the context of active transcription. Biochim. Biophys. Acta, Gene Regul. Mech. 2014; 1839,(12):1353–61.

5. Bannister AJ, Kouzarides T. Regulation of chromatin by histone modifications.Cell Res. 2011; 21, 381–395.

6. Olsen C. A. Expansion of the lysine acylation landscape. Angew. Chem., Int. Ed. 2012; 51, 3755–6.

7. Strahl BD, Allis CD. The language of covalent histone modifications. Nature 2000; 403, 41–45.

8. Wang YC, Suzanne E P and Loring JP Protein post-translational modifications and regulation of pluripotency in human stem cells. Cell Research 2014; 24: 143–160.

9. Wang Z, Yin H, Lau CS and Lu Q. Histone Post translational Modifications of CD4+T Cell in Autoimmune Diseases. Int. J. Mol. Sci 2016; 17: 1547.

10. Gillette TG, Hill JA. Readers, writers, and erasers: chromatin as the whiteboard of heart disease. Circulation research. 2015 Mar 27;116(7):1245–53.

11. Yang CH, Pospisilik AJ. Polyphenism –a window into gene-environment interactions and phenotypic plasticity. Frontiers in genetics. 2019; 10:132.

12. Mukherjee K, Twyman RM, Vilcinskas A. Insects as models to study the epigenetic basis of disease. Progress in biophysics and molecular biology. 2015; 118(1-2):69–78.

13. Kouzarides T. Histone methylation in Transcriptional control. Curr. Opin. Genet. Dev. 2002; 12, 198–209.

14. Fuhrmann J and Thompson P.R. Protein Arginine Methylation and Citrullination in Epigenetic Regulation. ACS Chem Biol. 2016; 11(3): 654–668.

15. Zhao Y and Garcia B A. Comprehensive catalog of currently documented histone meodification. Cold Spring Harb Perspect Biol. 2015; 7(9):a025064.

16. Kim J H, Yoo B C, Yang W S, Kim E, Hong S, and Cho JY. The Role of Protein Arginine Methyltransferases in Inflammatory Responses. Mediators of Inflammation. 2016; 1–11.

17. Kooistra S M, Helin K. Molecular mechanisms and potential functions of histone demethylases. Nat. Rev. Mol. Cell Biol. 2012; 13, 97–3119.

18. Black JC, Van RC, Whetstine JR. Histone lysine methylation dynamics: establishment, regulation, and biological impact. Mol Cell 2012; 48: 491–507.

19. Sims RJ 3rd, Nishioka K, Reinberg D. Histone lysine methylation: a signature for chromatin function. Trends Genet. 2003; 19: 629–39.

20. Shi Y and Whetstine JR. Dynamic Regulation of Histone Lysine Methylation by demethylases. Molecular Cell 2007; 25(1):1–14.

21. Cloos PAC, Christensen J, Agger K, and Helin K. Erasing the methyl mark: histone demethylases at the center of cellular differentiation and disease. Genes & Development 2008; 22, 1115–1140.

22. Tsai C, Shi Y and Tainer JA. How substrate specificity is imposed on a histone demethylase - lessons from KDM2A. Genes and Development (CSHP) 2014 28, 1735–1738.

23. Shi Y, Lan F, Matson C, Mulligan P, Whetstine JR, Cole PA, Casero RA, Shi Y. Histone demethylation mediated by the nuclear amine oxidase homolog LSD. Cell2004; 119, 941– 953.

24. Tsukada Y, Fang J, Erdjument-Bromage H, Warren ME, Borchers CH, Tempst P, Zhang Y. Histone demethylation by a family of JmjC domain-containing proteins. Nature 2006; 439, 811–816.

25. Kim T, Buratowski S. Two Saccharomyces cerevisiae JmjC domain proteins demethylate histone H3 Lys36 in transcribed regions to promote elongation. Journal of Biological Chemistry. 2007; 282(29):20827–35.

26. Paixão ES, Teixeira MG, Rodrigues LC. Zika, chikungunya and dengue: the causes and threats of new and re-emerging arboviral diseases BMJ Glob Health 2017; 3 e000530.

27. Gilbert C, Schaack S, Pace II JK, Brindley PJ, Feschotte C. A role for host-parasite interactions in the horizontal transfer of DNA transposons across animal phyla. Nature 2010; 464, 1347–50.

28. TheUniProtConsortium. UniProt: the universal protein knowledgebase Nucleic Acids Res. 2017; 45, D158–D169.

29. Emms DM, Kelly S. OrthoFinder: solving fundamental biases in whole genome comparisons dramatically improves orthogroup inference accuracy. Genome biology. 2015; 16(1):157.

30. Jones P, Binns D, Chang Hsin-Yu, Fraser M, Li W, McAnulla C, McWilliam H, Maslen J, Mitchell A, Nuka G, Pesseat S, Quinn AF, Sangrador-Vegas A, Scheremetjew M, Yong Siew-Yit, Lopez R, and Hunter S. InterProScan 5: genome-scale protein function classification Bioinformatics 2014; 30, 1236–4.

31. Katoh K, Standley DM. MAFFT multiple sequence alignment software version 7: improvements in performance and usability. Molecular Biology and Evolution 2013; 30:772–780.

32. Paradis E, Claude J and Strimmer K. APE: Analyses of phylogenetics and evolution in R language. Bioinformatics 2004; 20, 289–290.

33. Paradis E. Analyses of Phylogenetics and Evolution with R. Springer Science & Business Media; 2011.

34. Herrmann F, Pably P, Eckerich C, Bedford MT, Fackelmayer FO. Human protein arginine methyltransferases in vivo–distinct properties of eight canonical members of the PRMT family. J Cell Sci. 2009; 122(5):667–77.

35. Gaudet P, Livstone MS, Lewis SE and Thomas PD. Phylogenetic-based propagation of functional annotations within the Gene Ontology consortium. Briefings In Bioinformatics. 2010; 12, 449 – 462.

36. Stabell M, Larsson J, Reidunn BA, Lambertsson A. Drosophila dSet2 functions in H3-K36 methylation and is required for development. Biochemical and Biophysical Research Communications 2007; 359: 784–9.

37. Tsurumi A, Dutta P, Yan SJ, Shang R, Li WX. Drosophila Kdm4 demethylases in histone H3 lysine 9 demethylation and ecdysteroid signaling. Scientific reports. 2013 Oct 8; 3:2894.

38. International Aphid Genomics Consortium. Genome sequence of the pea aphid Acyrthosiphon pisum. PLoS biology. 2010 Feb 23;8(2):e1000313.

39. Elgin SCR and Reuter G. Position-Effect Variegation, Heterochromatin Formation and Gene Silencing in Drosophila. Cold Spring Harb. Perspect. Biol. 2013; 5, a017780.

40. Kang I, Choi Y, Jung S, Lim JY, Lee D, Gupta S, Moon W & Shin C. Identification of target genes regulated by the Drosophila histone methyltransferase Eggless reveals a role of Decapentaplegic in apoptotic signalling. Scientific Reports 2018; 8, 7123.

41. Clough E, Tedeschi T, Hazelrigg T. Epigenetic regulation of oogenesis and germ stem cell maintenance by the Drosophila histone methyltransferase Eggless/dSetDB1 Dev Biol 2014; 388, 181–191.

42. Rangan P, Malone CD, Navarro C, Newbold SP, Hayes PS, Sachidanandam R, Hannon GJ, Lehmann R. piRNA production requires heterochromatin formation in Drosophila Curr Biol 2011; 21, 1373–1379.

43. Smolko A, Shapiro-Kulnane L, Salz H. H3K9 methylation maintains female identity in Drosophila germ cells. Nature Communications 2018; 9(1):4155.

44. Yang SY, Chang YC, Wan YH, Whitworth C, Baxter EM, Primus S, Pi H, Van Doren M. Control of a novel spermatocyte-promoting factor by the male germline sex determination factor *Phf7* of *Drosophila melanogaster*. Genetics. 2017 Aug 1;206(4):1939–49.

45. Ogawa K, Miura T. Aphid polyphenisms: trans-generational developmental regulation through viviparity. Frontiers in physiology. 2014; 24:5:1.

46. Young LC, McDonald DW and Hendzel MJ. Kdm4b histone demethylase is a DNA damage response protein and confers a survival advantage following –irradiation. the journal of biological chemistry. 2013; 28, 21376–21388.

47. Colmenares SU, Swenson JM, Langley SA, Kennedy C, Costes SV, Karpen GH. Drosophila histone demethylase KDM4a has enzymatic and non-enzymatic roles in controlling heterochromatin integrity. Dev. Cell 2017; 42, 156–169.

48. Janssen A, Colmenares SU, Lee T, Karpen GH. Timely double-strand break repair and pathway choice in pericentromeric heterochromatin depend on the histone demethylase dKDM4A. Genes & Development. 2019; 33(1-2):103–15.

49. Vellichirammal NN, Gupta P, Hall TA, Brisson JA. Ecdysone signaling underlies the pea aphid transgenerational wing polyphenism. Proceedings of the National Academy of Sciences 2017; 114(6):1419–23.

50. Escandón-Vargas K, Muñoz-Zuluaga CA, Salazar L. Blood-feeding of Rhodnius prolixus, Biomédica, 2017; 37:299–302.

51. Kizer KO, Phatnani HP, Shibata Y, Hall H, Greenleaf AL, Strahl BD. A novel domain in Set2 mediates RNA polymerase II interaction and couples histone H3 K36 methylation with transcript elongation Mol Cell Biol 2005; 25(8):3305–16.

52. Brown DA, Di Cerbo V, Feldmann A, Ahn J, Ito S, Blackledge NP, Nakayama M, McClellan M, Dimitrova E, Turberfield AH, Long HK. The SET1 complex selects actively transcribed target genes via multivalent interaction with CpG island chromatin. Cell Reports 2017; 20 (10):2313–27.

