## Additional File 1 for "A comparative analysis of histone methyltransferases and demethylases in insect genome: A meta-analysis"

**A:** False positive rates in different insect proteome.

| **Class** | ***Drome*** | ***Aedae*** | ***Acypi*** | ***Rhopr*** | ***Apime*** | ***Bommo*** | ***Trica*** |
|---|---|---|---|---|---|---|---|
| Arginine methyltransferases | 0 | 0 | 0 | 0 | 0 | 0 | 0 |
| Lysine methyltransferases | 8 | 5 | 4 | 3 | 2 | 3 | 3 |
| Demethylases | 0 | 0 | 0 | 1 | 0 | 0 | 0 |

**B:** Novel proteins in different insect proteome

| **Class** | ***Drome*** | ***Aedae*** | ***Acypi*** | ***Rhopr*** | ***Apime*** | ***Bommo*** | ***Trica*** |
|---|---|---|---|---|---|---|---|
| Arginine methyltransferases | 0 | 0 | 4 | 4 | 2 | 4 | 0 |
| Lysine methyltransferases | 0 | 0 | 23 | 5 | 10 | 12 | 6 |
| Demethylases | 0 | 4 | 22 | 11 | 9 | 13 | 3 |

Drome: *Drosophila melanogaster*, Aedae: *Aedes aegypti*, Acypi: *Acyrthosiphon pisum*, Rhopr: *Rhodnius prolixus*, Apime: *Apis mellifera*, Bommo: *Bombyx mori*, Trica: *Tribolium castaneum*
