## Supplementary figures and images for "A comparative analysis of histone methyltransferases and demethylases in insect genome: A meta-analysis"

### Additional File 2

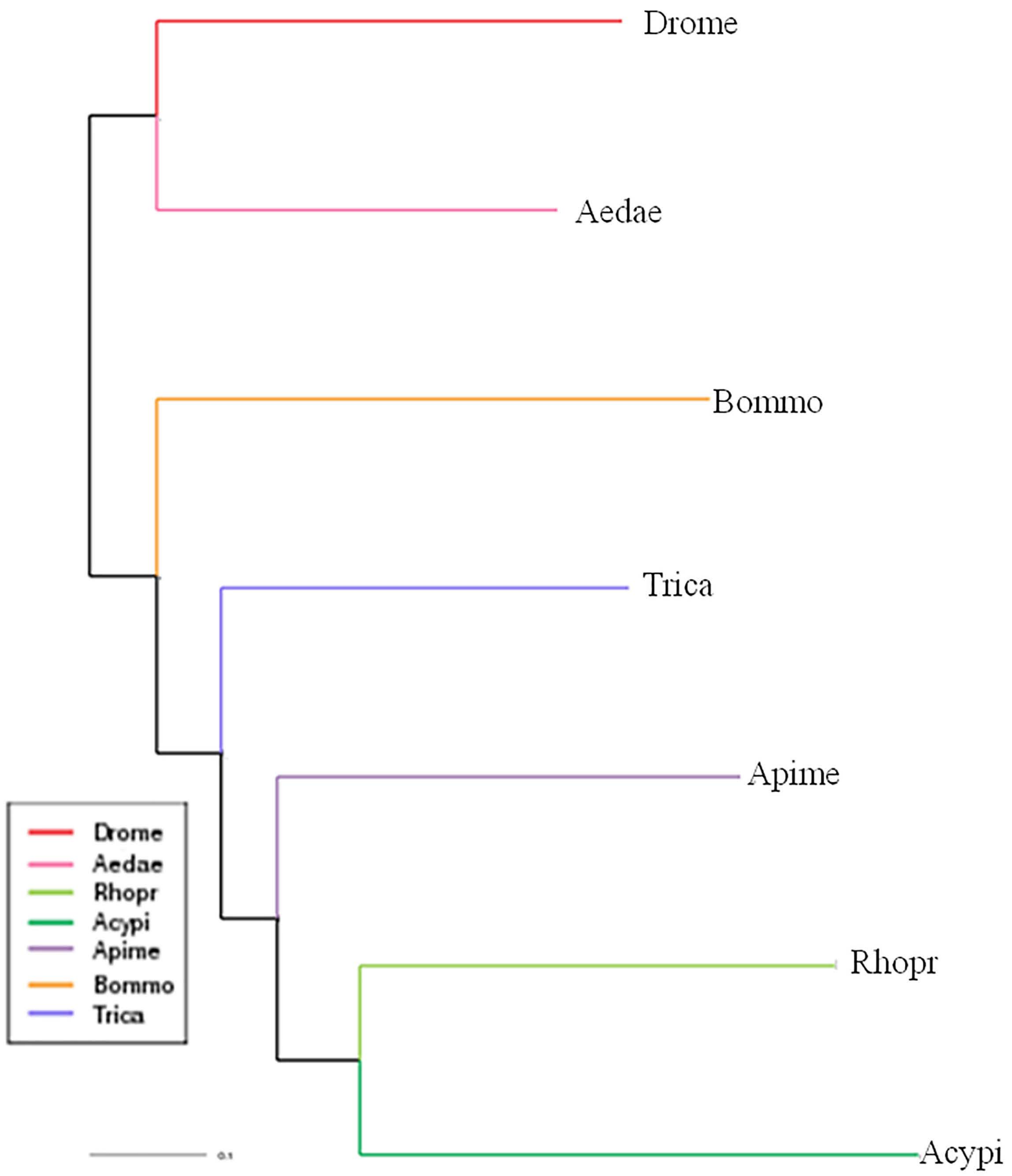
